# Quantifying Neural Stability: Validation of a New Brain Stability Index

**DOI:** 10.64898/2026.08.21.746247

**Authors:** Robert A. Seymour, Samuel Hardy, Yali Pan, Benjamin T. Dunkley

**Affiliations:** MYndspan, London, UK; Oxford Centre for Human Brain Activity, Department of Psychiatry, University of Oxford, Oxford, UK; Centre for Human Brain Health, School of Psychology, University of Birmingham, Birmingham, UK

**Keywords:** Magnetoencephalography, Brain Fingerprinting, Neural Stability, Synthetic Data, Neural Oscillations

## Abstract

Quantifying longitudinal changes in an individual’s brain is central to the development of personalised neural biomarkers in neurology and psychiatry. However, existing approaches for characterising individual neurophysiological signatures focus on discrimination between people rather than the quantification of within-subject change. To address this, we introduce the Brain Stability Index (BSI), a whole-brain metric that quantifies the similarity between two longitudinal neurophysiological scans in a low-dimensional latent space, with reference to a normative magnetoencephalography (MEG) database. Using 276 open resting-state MEG datasets and matched synthetic data, we first characterise how finite test-retest reliability sets a noise floor on the BSI. We then demonstrate that the BSI is sensitive to graded changes in whole-brain neural change that extend beyond measurement variability. Finally, we show that Factor Analysis, by separating shared structure from feature-specific noise, makes the BSI more robust to measurement artefacts. Together, these findings establish the BSI as a robust, bounded measure of neural stability that is well suited to longitudinal monitoring in neurology and psychiatry.

## 1 Introduction

The human brain exhibits an idiosyncratic functional architecture, with individual neurophysiological patterns increasingly recognised as possessing a stable, trait-like backbone that remains identifiable across recording sessions, cognitive tasks, and timescales (Ousdal et al. 2019; Ricchi et al. 2026; Dimitriadis et al. 2023). Because these patterns are stable within individuals, deviations from a subject’s own baseline can be used as a measure of neural change. Magnetoencephalography (MEG), with its high temporal resolution and spatial localisation (Baillet et al., 2017), is well suited to resolving these neurophysiological data features. Longitudinal MEG recordings have emerged as a promising route to individualised biomarkers of neurodevelopment, healthy aging (Gómez et al., 2013), and monitoring in psychiatry and neurology, including posttraumatic stress disorder (Zhang et al., 2020) and mild traumatic brain injury (Allen et al., 2020).

The dominant approach to individual MEG signatures has been brain fingerprinting, which demonstrates reliable discrimination between individuals even from short segments of data (Finn et al., 2015; da Silva Castanheira et al., 2021). Discrimination, however, is distinct from quantifying change within an individual. Identifiability establishes that subjects differ from one another (Amico et al., 2018) but provides no calibrated scale for the magnitude of within-subject change. Quantifying such change is demanding: MEG features are high-dimensional and sensitive to external artifacts as well as genuine neural change, and their finite test-retest reliability imposes a noise floor that any true effect must exceed to be detectable (Martín-Buro et al., 2016; Lew et al. 2021). Consequently, standardised, mathematically bounded metrics for single-subject longitudinal stability remain lacking.

To bridge this gap, we present the Brain Stability Index (BSI), a metric that quantifies the similarity between an individual’s longitudinal MEG scans in a low-dimensional latent space relative to a normative MEG database. Using simulations, we characterise its statistical properties and validate its ability to detect genuine neural change.

## 2 Materials and Methods

### 2.1 MEG Data Sources

To test the BSI, we sourced 276 openly-available MEG datasets and used these to generate additional synthetic data with matched univariate distributions and covariance structure. Resting-state MEG data were obtained from two open datasets:

i. *The Welsh Advanced Neuroimaging Database (WAND) (McNabb et al. 2025), comprising MEG, MRI, cognitive, and questionnaire data from 170 healthy adults aged 18–63. In total 162 participants had at least one MEG scan and generated usable processed MEG data*.
ii. *The Open MEG Archive (OMEGA) (Niso et al. 2016), an open dataset providing resting-state MEG, MRI, demographic, and disease data for healthy and Parkinsonian participants. 114 healthy participants were successfully processed and included*.

This yielded 276 resting-state MEG recordings in total. All participants provided informed written consent for sharing of anonymised data in the original studies. WAND was approved by the Cardiff University School of Psychology Research Ethics Committee (EC.18.08.14.5332RA3), and OMEGA access was approved by the Hospital for Sick Children Research Ethics Board under the OMEGA research agreement.

### 2.2 MEG Data Acquisition

MEG data in both WAND and OMEGA were acquired using whole-head 275-channel CTF systems while participants were in a magnetically shielded room. WAND recordings were collected at Cardiff University Brain Research Imaging Centre using a radial gradiometer configuration. OMEGA recordings were collected at the McConnell Brain Imaging Centre and Université de Montréal using an axial gradiometer CTF system. Both datasets obtained fiducial and head-shape information using a Polhemus digitisation system during participant preparation. Additionally, electrooculography and electrocardiography data were recorded alongside MEG. Resting-state recordings were acquired with participants seated upright with eyes open. WAND participants completed a 10-minute scan focussing on a white fixation dot to limit eye movements. Similarly, OMEGA resting-state recordings were eyes-open with a duration of at least five minutes, accompanied by a minimum of two minutes of empty-room data. MEG data was collected with a hardware low pass filter of 1200 Hz applied, and synthetic third-order gradiometry using reference magnetometers were used to reduce environmental magnetic interference.

### 2.3 MEG Data Processing

#### 2.3.1 Pre-processing

MEG data were processed using a custom analysis pipeline built on MNE-Python (Gramfort et al. 2013). The initial interference-reduction step depended on the acquisition system. For CTF recordings, environmental magnetic noise was attenuated using synthetic third-order gradiometry, which uses the CTF reference sensor array to estimate and suppress external magnetic interference (Vrba and Robinson 2001). For MEGIN/Elekta recordings, the pipeline first checked whether MaxFilter/signal space separation (SSS) had already been applied. If not, Maxwell filtering with temporal signal space separation (tSSS) was performed. When reliable continuous head position indicator (cHPI)-derived head-position information was available, head-motion compensation was incorporated into the Maxwell filtering step. Automated Maxwell-based bad-channel detection was also applied, allowing affected channels to be reconstructed during filtering.

Following system-specific interference reduction, all recordings were resampled to 300 Hz and band-pass filtered between 1 and 149 Hz. Power-line interference at either 50 or 60 Hz, depending on the country of data acquisition, together with its first harmonic, was removed using notch filtering. Physiological artefacts were removed using independent component analysis (ICA). For data that had not undergone Maxwell filtering, ICA was performed using 80 components identified for each scan using the infomax approach (Lee et al. 1999). For MaxFiltered data, the number of components was chosen to explain 98% of the variance. ICA components were classified using a proprietary machine-learning classifier, and components consistently identified as ocular or cardiac artefacts were excluded before reconstruction of the cleaned sensor-level data.

Finally, the cleaned recordings were segmented into non-overlapping 10.24 s epochs. Broadband artefacts were identified using a peak-to-peak amplitude threshold of 6000 fT, while muscle artefacts were detected from a 110–140 Hz filtered copy of the data using a peak-to-peak threshold of 2000 fT. Epochs overlapping either type of artefact annotation were rejected, and the remaining clean epochs were retained for subsequent source-space analyses.

#### 2.3.2 Source modelling

Magnetic lead fields were computed using a participant-specific single-shell boundary element model (BEM) derived from each participant’s anatomical MRI (Nolte 2003). T1-weighted MRIs were processed with FreeSurfer (Fischl 2012) to generate conformed anatomical images and Talairach transformations. Participant-specific inner-skull surfaces were obtained by transforming the fsaverage inner-skull surface into individual anatomical space and fitting a spherical harmonic representation, from which single-shell BEM solutions were computed.

Source modelling was performed using seventy-eight representative cortical locations derived from a reduced version of the Automated Anatomical Labeling (AAL) atlas, see Gong et al. (2009). All source locations were transformed into participant anatomical space to define a volumetric source space. A further 44 locations are included in the MYndspan pipeline for connectivity analyses, but data from these locations were discarded for this study. MEG-MRI co-registration was performed using the nasion and left and right preauricular fiducials to estimate the MEG-to-MRI transformation. Participant-specific forward solutions were then generated by combining a boundary element model (BEM) computed in MNE-Python, co-registration transform, and MEG sensor geometry.

Source reconstruction was performed using a linearly constrained minimum variance (LCMV) beamformer (Van Veen et al. 1997). LCMV beamforming is a spatial filtering method that reconstructs source activity by computing weighted linear combinations of the sensor-level MEG signals. The spatial filters are designed to pass activity originating from a specified source location with unit gain while minimizing contributions from all other sources. For each participant, cleaned sensor-level epochs were concatenated to estimate the data covariance matrix. Covariance matrices were regularized using Tikhonov regularization, with the regularization parameter set to 5% of the maximum singular value to improve numerical stability during matrix inversion. Spatial filters were computed at each of the predefined source locations using the participant-specific forward model, maximum-power orientation selection, and neural activity index (NAI) normalization to reduce spatial variations in noise sensitivity. Source activity was initially estimated using three orthogonal dipole orientations at each location, which were subsequently reduced to a single scalar source time course using singular value decomposition to determine the orientation of maximum power (Sekihara et al. 2004). The resulting source time courses were normalized to unit standard deviation.

Regional spectral power was estimated by computing the power spectral density (PSD) of the source time series extracted from the 78 AAL locations using Welch’s method with non-overlapping 4096-sample windows. Mean relative power was then computed within five predefined frequency bands: delta (1– 3 Hz), theta (4–8 Hz), alpha (9–12 Hz), beta (13–29 Hz), and gamma (30–45 Hz), with adjacent bands separated by 1 Hz to avoid overlap. This yielded a 390-dimensional feature vector (78 regions × 5 frequency bands) for each MEG recording, where each feature represented the relative spectral power of a specific brain region within a given frequency band. These feature vectors formed the input for the subsequent dimensionality reduction and brain similarity analyses.

### 2.4 Generating Synthetic Data

A Gaussian copula model (Nelsen 2006; Mai and Scherer 2017) was fitted to the combined WAND and OMEGA data feature matrix (276 recordings × 390 features, made up of 78 parcels × 5 frequency bands) using the copulas Python library. This model estimates the joint distribution of the data by separating the univariate marginal distribution of each feature from the dependency structure between features. Each feature was modelled as a beta distribution, allowing flexible estimation of marginal distributions, including for features with substantial skew.

Each MEG feature was transformed via its fitted marginal cumulative distribution function (CDF) to a uniform probability scale. These values were mapped into a latent Gaussian space where a single 390 × 390 covariance matrix captured the pairwise dependencies between all features. Synthetic observations were generated by sampling from this multivariate Gaussian distribution, then translated back into original MEG feature space using the inverse CDF of each marginal distribution. Because the shared covariance structure is Gaussian by construction, the low-dimensional linear structure subsequently recovered by Factor Analysis is present by design rather than discovered empirically; the synthetic data should therefore be read as a controlled test of BSI behaviour under known reliability and known change, not as independent evidence about the true dependency structure of MEG power.

This method was used as the MEG feature vector contains variables with different ranges and distributions. By fitting flexible marginal distributions independently for each feature the underlying univariate distributions can be preserved while approximating the observed multivariate structure. Once fitted, the model was sampled 1000 times, producing a synthetic dataset of MEG-like observations with the same dimensionality, approximate univariate distributions and covariance structure as real MEG datasets.

## 3 Results

### 3.1 The Brain Stability Index (BSI)

We introduce a novel metric to quantify the similarity between two MEG scans, with reference to a normative database of resting-state MEG data. The metric is designed to be used longitudinally, tracking data features from the same individual across repeated datasets. Each scan is first represented as a 390-dimensional feature vector comprising band-limited spectral power (78 parcels × 5 frequency bands, see Methods). To reduce dimensionality and isolate shared structure across participants, we fit a Factor Analysis (FA) model (45 factors, quartimax rotation) to a normative resting-state sample (see Methods). The number of factors was determined empirically using cross-validation (Supplementary Figure 1). Each dataset was projected into this shared factor space, yielding a 45-dimensional factor score vector, z.

The Brain Stability Index (BSI) between two scans from the same individual is then calculated as:

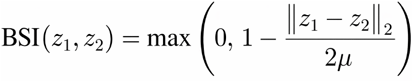

where ‖z1 − z2‖ is the Euclidean distance between the two scans in factor space, and μ is the mean pairwise Euclidean distance across between-subject pairs in the same normative sample of resting-state MEG data. The role of μ is to anchor the scale: BSI = 0.5 corresponds exactly to the average dissimilarity between two randomly selected individuals in the normative sample (Figure 1). Values are clipped at 0 so the index lies in [0, 1]. Values above 0.5 indicate that two scans are more alike than a typical pair of different people, while values below 0.5 indicate divergence beyond normal inter-individual variation. A BSI of 0 corresponds to scans differing by at least twice the mean between-person distance, a degree of change rarely seen in longitudinal MEG data (see Table 1).

**Table 1.**
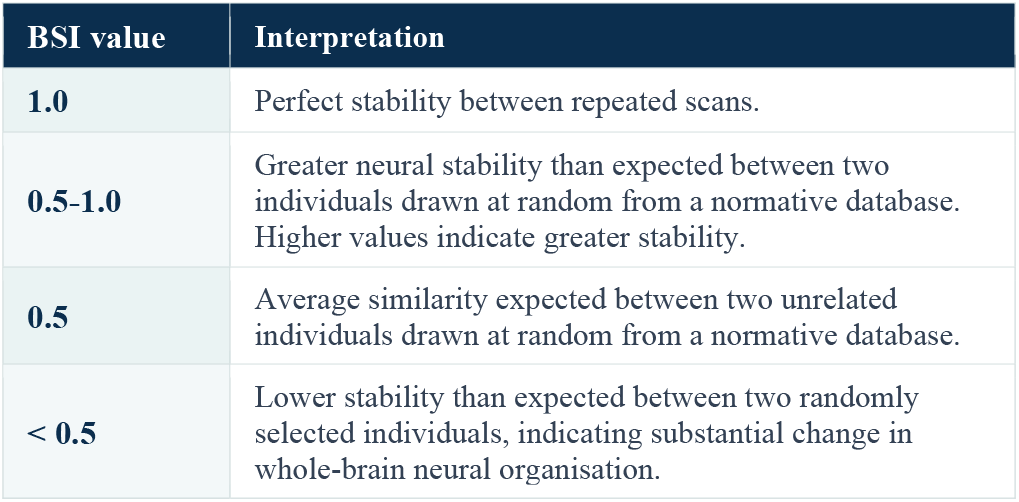
Interpretation of Brain Stability values.

**Figure 1.**
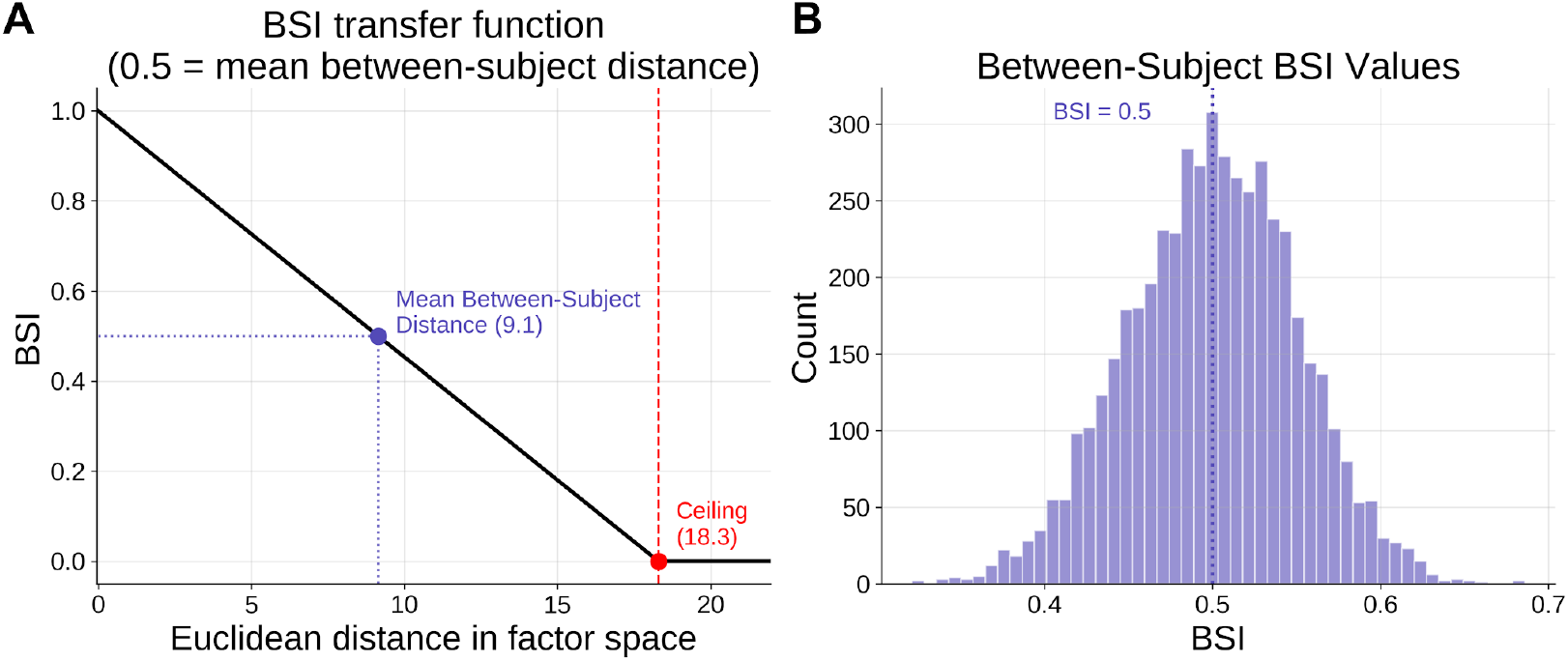
Brain Stability Index (BSI) scaling and between-subject distribution. (A) The BSI transfer function converts the Euclidean distance between MEG features in factor space to a 0–1 similarity score, anchored so that the mean between-subject distance equals BSI = 0.5 and distances beyond twice this value are clipped to zero. (B) Between-subject BSI values from simulated data approximate a normal distribution centred on 0.5.

### 3.2 Validating BSI

To validate the BSI we generated 1000 synthetic 390-dimensional feature vectors with the same statistical properties as real MEG resting-state data (see Methods), and anchored the BSI scale (μ) using the mean distance across all 499,500 unique between-subject pairs. An important consideration for any longitudinal metric is the reliability of the underlying measurement. MEG features have finite test–retest reliability, typically quantified by the intraclass correlation coefficient (ICC). This means that even when no true brain change has occurred, repeated scans from the same individual will not produce identical feature vectors: measurement noise introduces a floor on the minimum detectable change. We therefore characterised how BSI behaves under known levels of ICC, so that longitudinal changes can be distinguished from measurement variability. Synthetic retest scans were generated by adding Gaussian jitter to each factor score, with the jitter scaled so that the resulting test-retest reliability matched a target ICC. As expected, we found that as ICC decreases, BSI also decreases (Figure 2A). This relationship is nonlinear because within-subject variance scales as (1 − ICC) / ICC: noise grows slowly as ICC drops from 1.0 but accelerates rapidly below ~0.4. At an ICC of 0.7, typical of alpha- and beta-band spectral power in MEG (Lew et al., 2021), the retest BSI was 0.77 (95% CI: 0.71–0.82), meaning that approximately 23% of the BSI scale is consumed by measurement noise.

**Figure 2.**
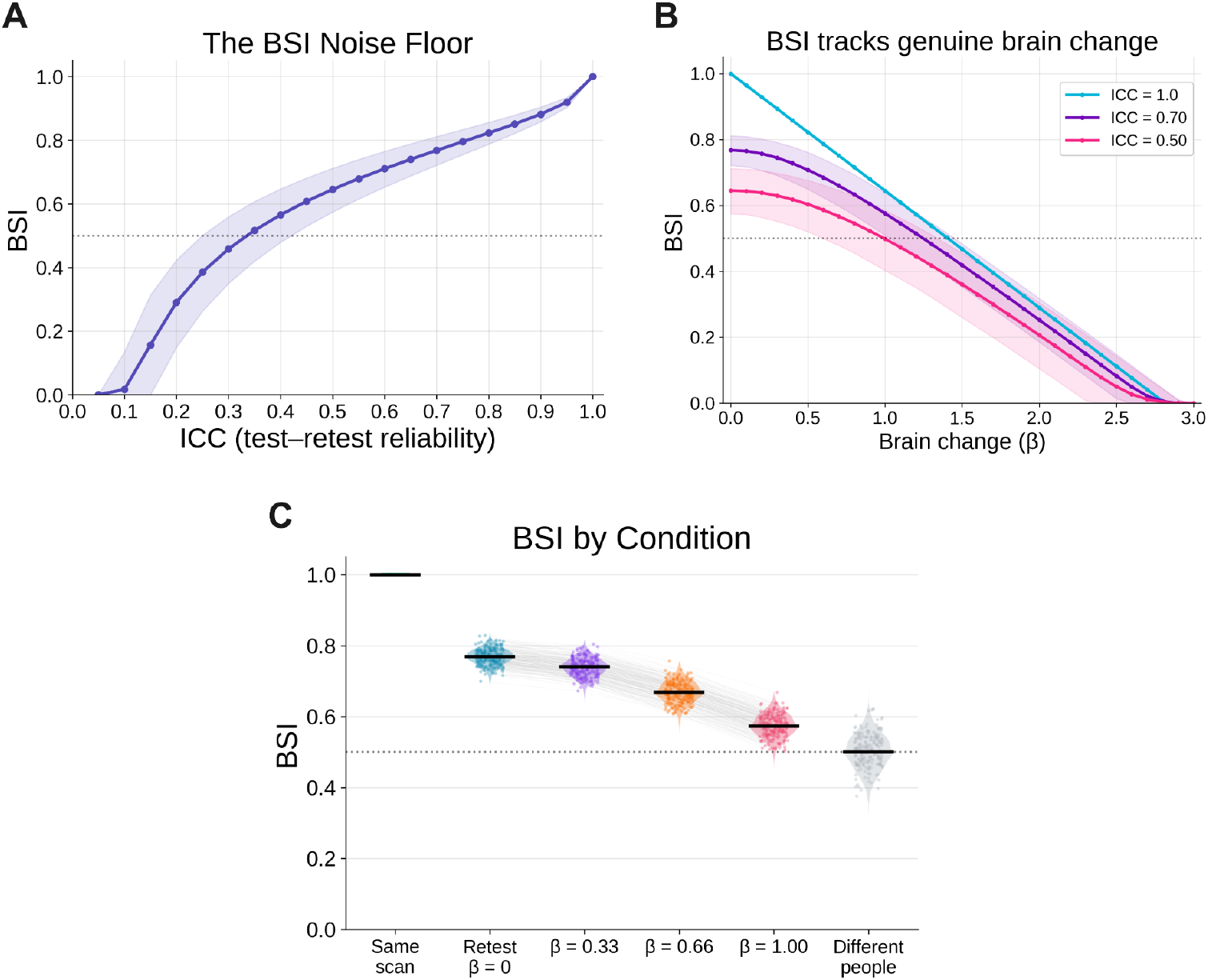
(A) BSI plotted as a function of test-retest reliability (ICC) to establish a noise floor. (B) BSI plotted as a function of genuine brain change (β, between-subject SDs in factor space) at three ICC levels. Across plots, the dotted line at BSI = 0.5 marks the mean between-subject distance: change beyond this point looks as large as the difference between two random people. Shaded bands, 95% CI across 1000 synthetic participants. (C) BSI distributions for six conditions at ICC = 0.7: same scan, retest with no brain change (β = 0), graded simulated brain change (β = 0.33, 0.66, 1.0), and different participants. Black bars indicate the median; grey lines track individual participants across change levels.

To evaluate BSI sensitivity to genuine longitudinal brain change above the noise floor, we simulated shifts in each dataset’s feature vector toward another randomly selected feature vector of the 1000 datasets. Shift magnitude was parameterized by β, measured in units of the between-subject standard deviation in factor space, where β = 0 denotes no change and β = 1.0 denotes a one-standard-deviation shift. As expected, BSI decreased with the magnitude of change (Figure 2B). At ICC = 1.0, this relationship is perfectly linear. At ICC = 0.7 and ICC = 0.5, the entire curve shifts downward by the noise floor but retains a negative slope. This shows that realistic measurement noise lowers the BSI ceiling without distorting overall sensitivity to change.

Finally, to provide a practical reference for interpreting BSI in real data, we fixed ICC at 0.7 — a value representative of alpha- and beta-band spectral power in MEG (Martín-Buro et al. 2016; Lew et al. 2021). For each of the 1000 synthetic datasets, we plotted the BSI score given no neural change (retest; β = 0). Then for each dataset we combined measurement variability with genuine brain change by shifting each dataset’s factor score vector in factor space toward a randomly selected dataset, at β = 0.33, 0.66 and 1.00 (same units as above). We also plotted the BSI calculated against exactly the same scan (no measurement noise, BSI = 1) and the between-dataset BSI scores for comparison. BSI differed across all six conditions (Figure 2C). Critically, even a β = 0.33 shift was distinguishable from β = 0 (i.e. no neural change), and a β = 1.00 shift approached the between-subject distribution. This shows that given a fixed level of measurement reliability (as would be expected in real MEG data) BSI provides graded sensitivity to neural change within an interpretable, bounded scale.

### 3.3 Investigating Factor Analysis

Factor Analysis (FA) was selected as the dimensionality reduction step in the BSI pipeline because, unlike alternative methods such as principal component analysis, it explicitly models per-feature measurement noise separately from shared variance. We hypothesized that FA would improve the robustness of the BSI to measurement artefacts distributed across data features (e.g., arising from head movement, sensor drift, or environmental interference). To test this, we compared two pipelines: one in which BSI was computed in a 45-dimensional factor space (with FA), and another in which BSI was computed in the full 390-dimensional feature space (without FA). Both pipelines were calibrated such that BSI = 0.5 corresponded to the mean between-subject distance in their respective feature spaces.

## 4 Discussion

Accurate measurement of longitudinal brain change at the single-subject level is fundamental to personalised, data-driven healthcare across neurology and psychiatry (Uhlhaas et al. 2017). However, the high dimensionality and intrinsic variability of neurophysiological data have made robust, objective quantification of longitudinal brain stability challenging. In this study, we introduce and comprehensively validate the Brain Stability Index (BSI), demonstrating its utility as a standardised, noise-resilient metric for quantifying longitudinal resting-state MEG data. By anchoring the BSI against a large normative MEG dataset and scaling the metric between 0 and 1, the BSI provides a directly interpretable measure of neural stability (Table 1 and Supplementary Table 1 for lay interpretations).

We applied the same broadband measurement noise to all 1,000 synthetic participants at graded intensities (α = 0 to 2.5, where α scales each feature’s noise relative to its between-subject standard deviation; α = 1 means measurement noise equal in magnitude to natural between-person variation on each feature). As shown in Figure 3, when using FA, BSI dropped less steeply than the BSI without FA, confirming that FA makes the metric more robust to measurement noise. This can also be seen when comparing BSI values from paired synthetic datasets - with and without FA (Supplementary Figure 2A).

**Figure 3.**
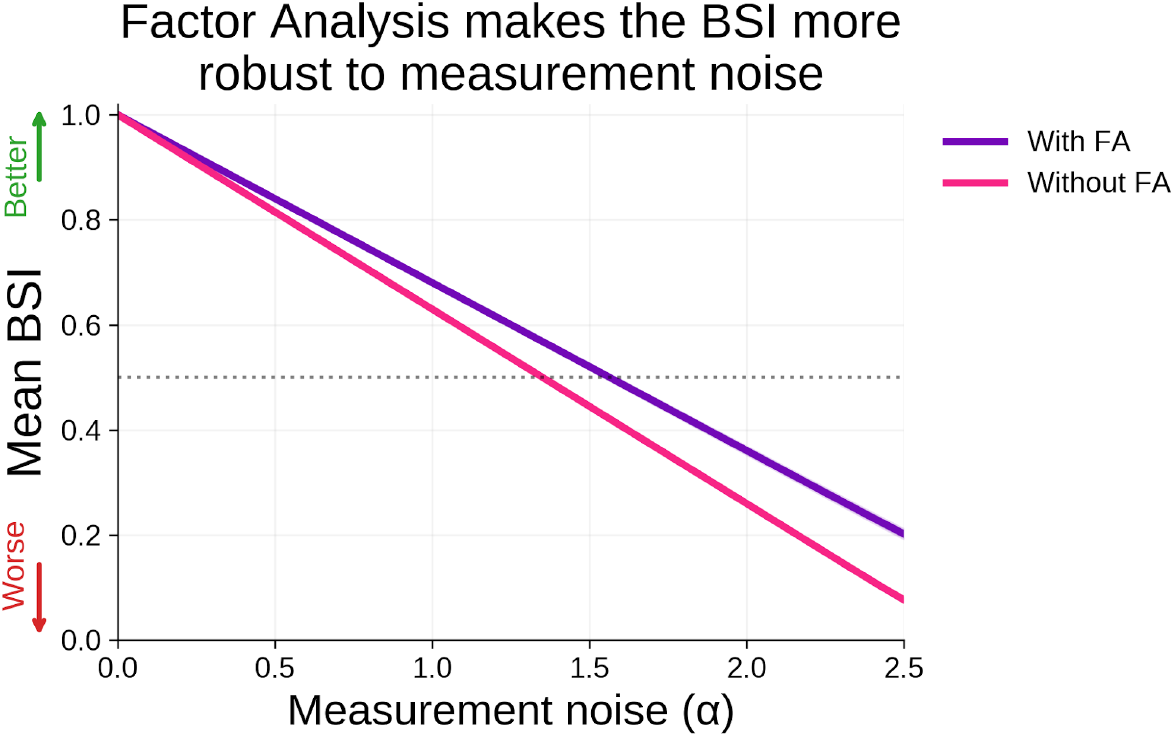
BSI calculated using a Factor Analysis (FA) step and without FA, with both versions anchored such that BSI = 0.5 at the mean between-subject distance (dotted line). Measurement noise was then added to each synthetic dataset at varying levels (α), and BSI was computed for the FA (purple) and no-FA (pink) conditions.

We also repeated the analysis using raw Euclidean distances to assess whether the apparent FA advantage was driven by differences in BSI scaling. As shown in Supplementary Figure 2B, this was not the case: incorporating FA into the pipeline caused the noise-induced Euclidean distance to grow more slowly with increasing measurement noise (α), even after normalising by between-subject spread in each space.

In this technical report we show that the BSI is a robust and reliable marker of longitudinal brain change. Using synthetic data we were able to explicitly model the effect of finite test-retest reliability on BSI scores (Uhlhaas et al. 2017). This builds on prior work showing that spectral power in the alpha and beta bands are highly robust, frequently exhibiting ICCs > 0.60 (Lew et al. 2021). Establishing this noise floor (Figure 2A) is an important empirical step for BSI, because it defines how much a score can vary between recordings of an unchanged brain, and therefore the threshold a real change must exceed to be distinguishable from measurement error. Even though the BSI is practically constrained by imperfect measurement reliability, using simulated data we showed that the BSI retains graded sensitivity to shifts in neural stability (Figure 2B): larger underlying changes produce proportionally larger BSI shifts. Importantly, given a fixed test-retest reliability of 0.7 reminiscent of real MEG data, simulated shifts in neural change rose clearly above the noise floor and were reliably detectable using the BSI (Figure 3).

A further innovation in the BSI pipeline is the use of factor analysis (FA) for dimensionality reduction. In simulations using a large battery of existing MEG-derived features, our comparative analysis shows that computing similarity in lower-dimensional latent factor space outperforms the use of the full set of 390 power features. FA separates shared covariance structure from feature-specific variance, such that noise arising from environmental interference, sensor drift, or other measurement variability does not consistently load onto latent dimensions and is therefore attenuated. The resulting BSI is more robust to measurement noise, leveraging shared structure across existing MEG data to isolate stable neural signals (Seymour et al., 2022) across individuals and sessions.

More broadly, there are substantial clinical applications for measures of whole-brain neural stability like the BSI, particularly for longitudinal monitoring and therapeutic evaluation. Many neurological disorders are characterised by progressive, transient, or recovery-related changes in large-scale brain networks that are difficult to quantify in individual patients. These include the evolving phases of mild traumatic brain injury (Vakorin et al. 2016; Edgar et al. 2026), neural reorganisation following stroke (Santoro et al. 2026), and the early stages of Alzheimer’s disease (Stampacchia et al. 2024) and Parkinson’s disease (Castanheira et al. 2024). Across these conditions, the Brain Stability Index (BSI) could quantify longitudinal changes in whole-brain neural stability, offering a quantitative marker of disease progression, recovery, or treatment response. The BSI may also complement existing clinical and behavioural outcomes and serve as a quantitative endpoint in clinical trials (Lanskey et al. 2022). This could improve the evaluation of pharmacological and neurorehabilitative interventions by providing an objective measure of changes in whole-brain network organisation.

Despite these promising results, three limitations should be noted. First, the current validation relies on synthetic data generated using copula modelling. Although this approach preserves the distributions and covariance structure of the empirical MEG data, synthetic modelling depends on the assumption of multivariate normality within latent space. Future empirical work should therefore validate the BSI using real, multi-session longitudinal datasets acquired across multiple scanning sessions and scanner types (Larson-Prior et al., 2013), as well as in contexts where neural change is likely to occur, such as mild traumatic brain injury (mTBI), stroke, or following psychiatric episodes. This would help establish a normative basis for BSI values and improve our understanding of longitudinal brain change more broadly. Second, we note that the current formulation of the BSI operates on oscillatory power alone. Future work is planned to explicitly combine power with connectivity data (e.g. amplitude-envelope correlations), while applying dimensionality reduction techniques to equate data features. Third, the BSI is a metric that averages MEG data features over time and is therefore highly suited to resting-state data. However, this temporal averaging obscures fast fluctuations in brain dynamics that occur at shorter timescales. Future work could address this limitation by explicitly modelling transient brain states using approaches such as microstate analysis or Hidden Markov Models (HMMs), which capture rapid state transitions in electrophysiological data (Michel & Koenig, 2018; Baker et al., 2014; Vidaurre et al., 2018). This would allow derivation of a time-resolved marker of brain stability during ongoing cognition across a range of experimental paradigms.

In conclusion, the Brain Stability Index represents a robust and interpretable measure of neural stability for use in longitudinal neurophysiological datasets.

## AUTHOR CONTRIBUTIONS

**RAS:** Conceptualization, Methodology, Formal analysis, Investigation, Visualization, Writing - original draft, Writing - review & editing.

**SH:** Conceptualization, Methodology, Data curation, Formal analysis, Visualization, Writing - original draft, Writing - review & editing.

**YP:** Conceptualization, Methodology, Writing - original draft, Writing - review & editing.

**BTD:** Conceptualization, Methodology, Project administration, Supervision, Writing — original draft, Writing - review & editing.

## ACKNOWLEDGMENTS

We acknowledge Dr. Gill Roberts who conducted some initial pilot analyses that led to the computation of the Brain Stability Index.

## DATA AVAILABILITY

The datasets analysed for this study are openly available from their original repositories. The Welsh Advanced Neuroimaging Database (WAND) is available through the G-Node GIN repository (https://doi.gin.g-node.org/10.12751/g-node.5mv3bf/; McNabb et al., 2025). The Open MEG Archive (OMEGA) is available at https://omega.bic.mni.mcgill.ca (Niso et al., 2016).

## SUPPLEMENTARY MATERIAL

**Supplementary Figure 1.**
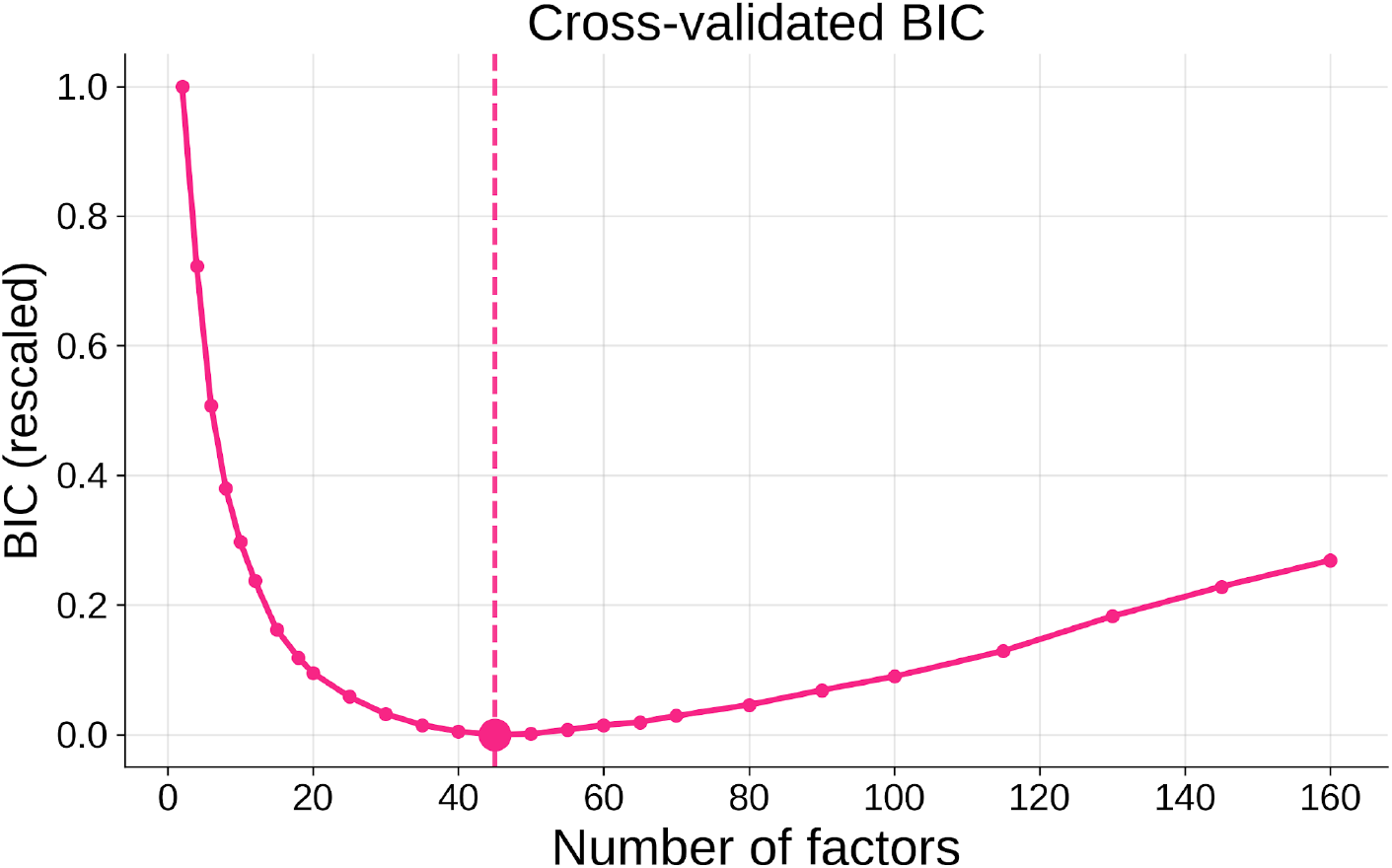
Determining the number of BSI factors via cross-validated BIC. Factor Analysis models were fit to the 390-dimensional power feature block (78 cortical ROIs × 5 frequency bands) across a range of factor counts (2–150). For each iteration, the 1000 synthetic datasets were partitioned into 5 folds. A Factor Analysis model was fit on 4 folds and assessed on the held-out fold, repeated so that every participant was scored exactly once out-of-sample. For each iteration, held-out log-likelihood was summed across folds and converted to the Bayesian Information Criterion (BIC) to penalise complexity as the number of factors rises. BIC is plotted as ΔBIC scaled between 0-1. The lowest value was present at 45 factors (pink marker, dashed line).

**Supplementary Figure 2.**
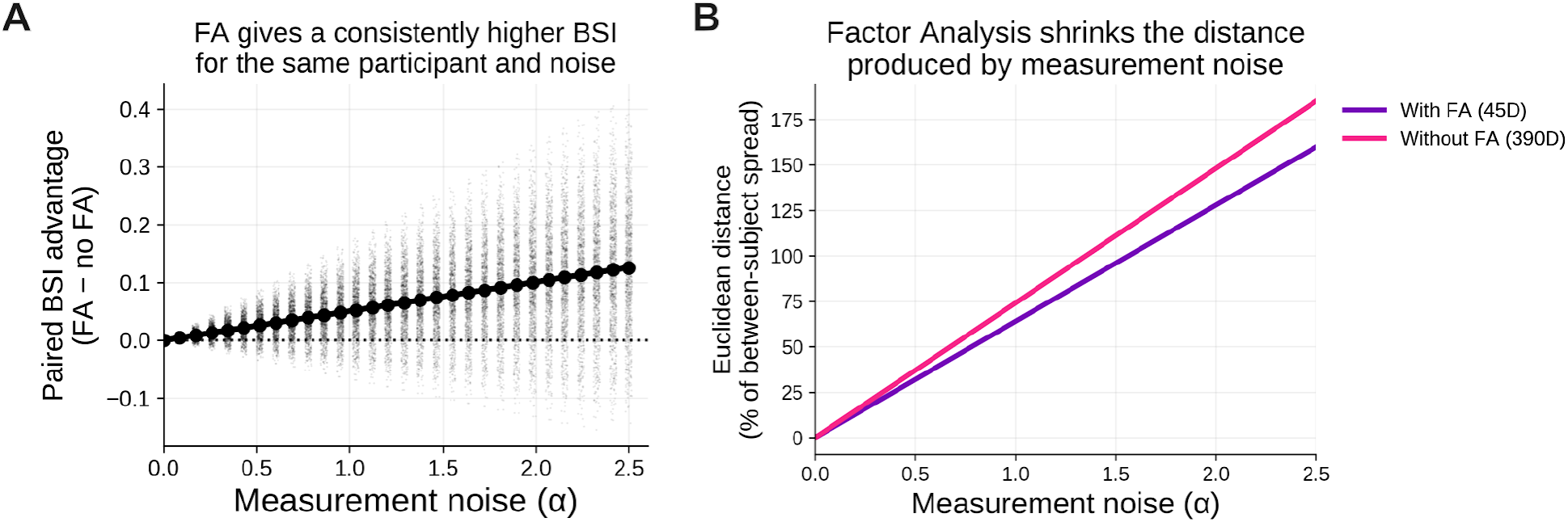
(A) For each synthetic dataset at each noise level, BSI was computed twice (once with the factor analysis step included and once with it omitted) and the difference between the two scores was calculated. While there is some variation, there is a clear advantage when including factor analysis, especially at higher measurement noise levels. (B) Rather than compute the BSI as measurement noise increases, we instead calculated raw Euclidean distance, normalised as a % change versus the between-subject spread. Again we see a clear protective effect when including factor analysis - the Euclidean distance grows less with measurement noise.

**Supplementary Table 1.** Lay interpretation of Brain Stability values.

| BSI | Plain label | Lay interpretation |
| --- | --- | --- |
| $\geq 0.75$ | Stable | Your brain pattern is highly consistent with your previous scan, showing only small differences. |
| $0.5 - 0.75$ | Some change | This is within the normal range of variation. Your brain has changed since your last scan, but the amount of change is within the range typically seen between scans in healthy individuals. |
| $< 0.5$ | Large change | There has been a substantial change since your last scan. The similarity between scans is comparable to the average level of similarity expected between two unrelated people. |

